# svviz: a read viewer for validating structural variants

**DOI:** 10.1101/016063

**Authors:** Noah Spies, Justin M. Zook, Marc Salit, Arend Sidow

## Abstract

Visualizing read alignments is the most effective way to validate candidate SVs with existing data. We present svviz, a sequencing read visualizer for structural variants (SVs) that sorts and displays only reads relevant to a candidate SV. svviz works by searching input bam(s) for potentially relevant reads, realigning them against the inferred sequence of the putative variant allele as well as the reference allele, and identifying reads that match one allele better than the other. Reads are assigned to the proper allele based on alignment score, read pair orientation and insert size. Separate views of the two alleles are then displayed in a scrollable web browser view, enabling a more intuitive visualization of each allele, compared to the single reference genome-based view common to most current read browsers. The web view facilitates examining the evidence for or against a putative variant, estimating zygosity, visualizing affected genomic annotations, and manual refinement of breakpoints. An optional command-line-only interface allows summary statistics and graphics to be exported directly to standard graphics file formats. svviz is open source and freely available from github, and requires as input only structural variant coordinates (called using any other software package), reads in bam format, and a reference genome. Reads from any high-throughput sequencing platform are supported, including Illumina short-read, mate-pair, synthetic long-read (assembled), Pacific Biosciences, and Oxford Nanopore. svviz is open source and freely available from https://github.com/svviz/svviz.

## Introduction

The human eye has an unparalleled ability to identify patterns from visual representations of data. While the identification of mutations from high-throughput sequencing has been largely automated, visual inspection of putative variants using tools such as the Integrative Genomics Viewer (Robinson 2011) remains an important step in ensuring the quality and relevance of these variant calls. However, existing read visualizing tools such as IGV are largely constrained by a reference-genome-centric display model. Hence, point mutations can be represented easily as mismatched bases within sequencing data, but more complex structural variants (SVs) including insertions, deletions, translocations and inversions are more difficult to parse visually against the linear reference genome sequence. Newer tools, such as Bambino (Edmonson 2011) and PyBamView (Gymrek 2014), are able to represent short indels within sequencing data but do not help in representing larger SVs.

Support for SVs can be displayed within IGV by highlighting reads with certain characteristics, including read pairs mapping to distant regions of the genome or in unexpected orientations, or truncated alignments. However, it is difficult to identify from these highlighted, discordantly mapping reads whether they all agree with a putative variant, and if so, which variant. Furthermore, IGV relies on the quality and completeness of the alignments provided in input BAM files, which are produced en masse against a huge reference genome and hence may not optimally represent read support for a given variant. Finally, most existing viewers (a notable exception being TargetSeqView; Halper-Stromberg 2014) show all read data in the vicinity of a putative structural variant, making it difficult to discriminate reads supporting the SV, reads supporting the reference allele, and reads that are not relevant to an SV. This can be nearly impossible when physical coverage is extremely high, for example when looking at mate-pair data.

To overcome these limitations, we present svviz, a read visualizer for structural variants that sorts and displays only reads relevant to the current SV. As with IGV, svviz only visualizes variants and does not identify them. svviz runs locally on a standard OS X or linux desktop machine, and requires as input read data, a reference genome, and structural variants. The flexible approach employed by svviz means it can display arbitrary SV types such as deletions, insertions, inversions and mobile element insertions (support for translocations and other more complex events is planned but not yet implemented). Visualizations are rendered in SVG (scalable vector graphics), an open web standard graphics format, and shown in a locally-hosted interactive web browser viewer or exported in publication-ready form. svviz supports read data in BAM format from any sequencing platform, including short-read [Illumina (Bentley 2008)] single-and paired-end as well as mate-pair or longer read [Pacific Biosciences (Eid 2009), Oxford Nanopore, or Illumina’s synthetic long-reads] sequencing technologies.

## Methods

svviz performs several pre-processing steps before visualizing a particular structural variant:

1. Breakpoints for the input SV are processed to produce a representation of the unique genomic sequence of the SV.
2. Reads are identified that map near all SV breakpoints.
3. For paired end data, read mates are collected.
4. Reads are realigned to both the alternate (SV) allele, as well as the reference allele, by Smith-Waterman alignment (Zhao 2013).
5. Read pairs (or reads, when run in single-ended mode) are assigned to reference or alternate alleles if they better support one allele over the other, or are otherwise labeled as ambiguous. The criteria for this step are described below.
6. Reference and alternate alleles are visualized separately with individual tracks for each input sample for each allele (especially useful for related samples in a family pedigree). Ambiguous reads, typically mapping near but outside of the breakpoints, can also be visualized in a third set of tracks, aligned against the reference sequence.

Realigned reads are assigned to the allele with the higher alignment score, or (if scores are identical), the allele with the better match to the empirical insert size distribution (derived from the input BAM file). Only allele assignments with alignment scores exceeding a threshold (adjustable based on the error rates of the sequencing technology), and with the expected read pair orientation(s) are considered. Reads (or read pairs) that cannot confidently be assigned to one allele or the other are instead marked as ambiguous.

This process extends the approach adopted in TargetSeqView (Halper-Stromberg 2014), enabling allele assignment of reads of arbitrary length (as are typically produced by the long-read technologies) and taking advantage of the insert size distribution, which can be more informative than the alignment score. In addition, use of the empirical insert size distribution provides a more accurate discrimination between alleles than the z-score-based approach typically adopted by SV calling software (eg Delly; Rausch 2012).

## Results

The Genome in a Bottle consortium (Zook 2014) has recently begun sequencing an Ashkenazi Jewish trio from the Personal Genome Project with a number of high-throughput sequencing platforms, providing a rich resource for identifying and validating variants using orthogonal experimental methods. Structural variants were called from Complete Genomics data for mother, father and son separately. From these variant calls, we randomly chose an 11.5kb inversion on chromosome 4 to visualize and validate using long mate-pair Illumina data.

The visual representation is split into two sections. The top section (Figure 1a) shows reads supporting the alternate “inversion” allele, while the bottom section (Figure 1b) shows those supporting the reference allele. Each read is shown only once, shown relative to its assigned allele. The axis below shows the inverted region as blue with arrows pointing to the right in the reference allele and to the left in the alternate allele. Adjacent genomic regions are colored red and gray. The first and second read ends from these long insert-size mate-pair data are shown as red and purple boxes, connected by light gray lines.

**Figure 1.**
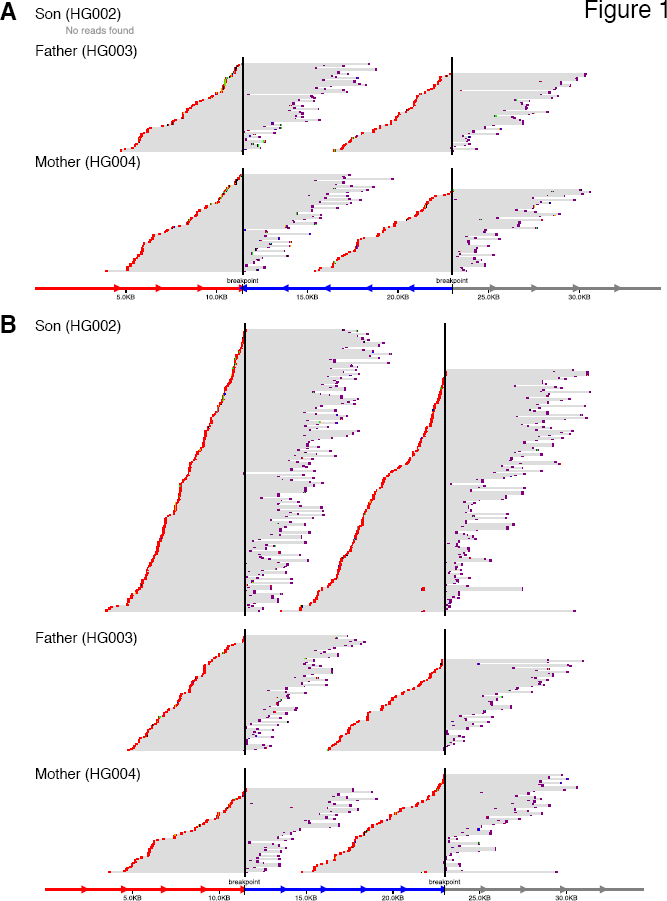
svviz visualization of chr 4 inversion. (a) Reads supporting the alternate allele, with inverted region represented as blue region at bottom, between marked breakpoints. Blue arrows pointing to the left demarcate the inverted region. (b) Reads supporting the reference allele, with non-inverted region again shown in blue at bottom. Blue arrows in between breakpoints point to the right, indicating all genomic sequences in the reference allele are in the same orientation. Red reads are on the minus strand and purple reads are on the plus strand, with gray lines linking mate-pairs (note that the mate-pair data shown here are sequenced in -/+ orientation, and have an average insert size of ~6.5kb). Ambiguous reads, those unable to distinguish between the alleles, are not shown.

For the alternate allele, mate-pair reads tile across the breakpoints in both parents, although no reads were found in the son. All three individuals show ample coverage of the reference allele breakpoints. The number of reads assigned by svviz to each allele suggests the son is homozygous reference and both parents are heterozygous for the inversion. Figure S1 shows the same data represented in IGV, with likely non-reference reads colored maroon and blue, suggestive of a structural variant but difficult to interpret as an inversion. Figure S2 shows a putative 1200bp deletion for which svviz shows very little supporting evidence, and which we estimate is a false-positive.

svviz can typically be installed on OS X and linux using the single command “`sudo pip install svviz`”. Documentation is available at http://svviz.readthedocs.org and the source code is available from https://www.github.com/svviz/svviz. Please report any difficulties installing or running svviz on the github issues page.

The inversion shown in Figure 1 is included as example data with svviz, and can be downloaded and visualized in the interactive web browser view simply by running the command “`svviz demo`” from the command-line.

## Acknowledgments

We thank the Genome in a Bottle Consortium and Complete Genomics for making the data publicly available. Certain commercial equipment, instruments, or materials are identified in this document. Such identification does not imply recommendation or endorsement by the National Institute of Standards and Technology, nor does it imply that the products identified are necessarily the best available for the purpose.

## Figure Legends

**Figure S1.**
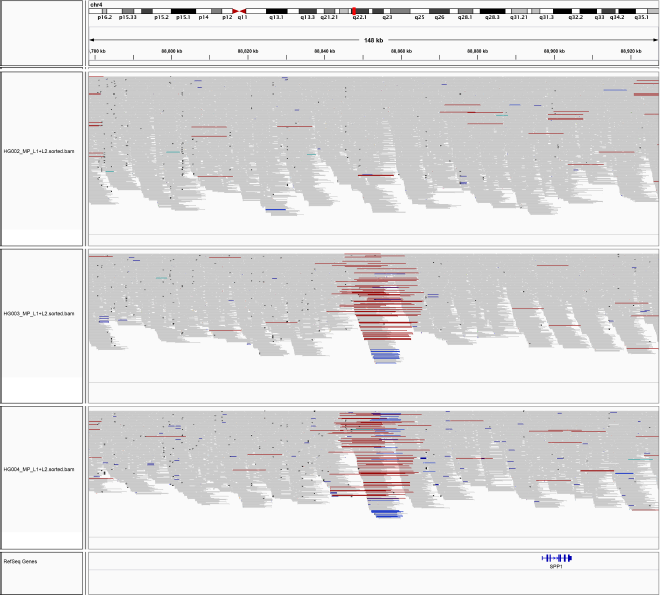
Same chromosome 4 inversion as in Figure 1, visualized in IGV. Maroon and blue reads align to the genome in incorrect orientation or unexpected insert-size.

**Figure S2.**
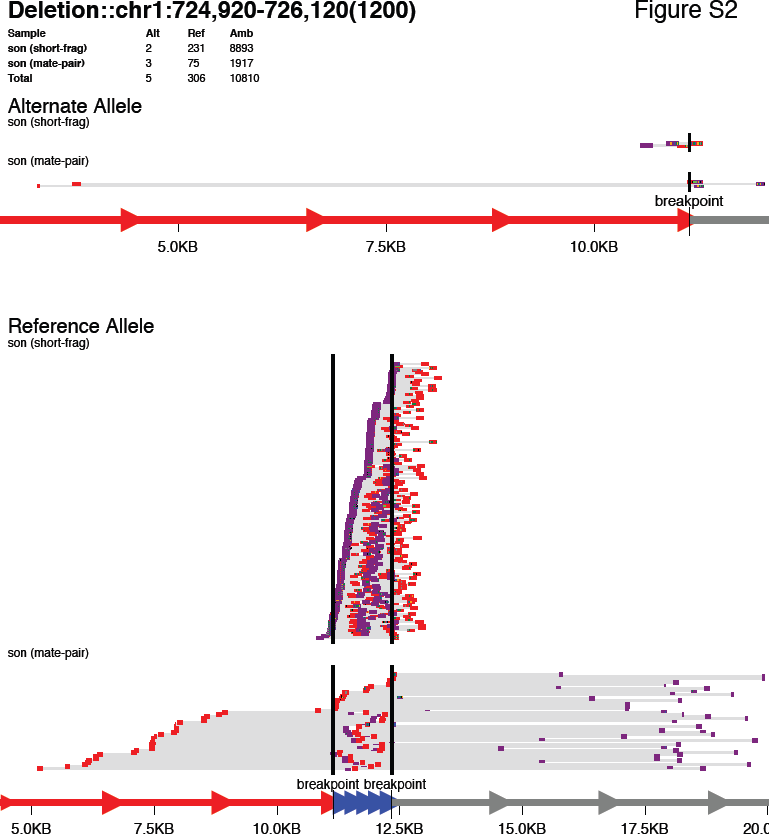
Example raw output from svviz of a false-positive deletion called in the son of the Ashkenazi trio. Several features demonstrate this is unlikely to be a true deletion. (1) The overall coverage is extremely high (see the overview table at the top, showing high numbers of ambiguous reads, suggesting the region is likely repetitive). (2) Only a small number of reads cover the breakpoints in the Illumina short-fragment library (top track) and in the mate-pair library (bottom track). Furthermore, it is clear in the web view that these reads have many mismatches to the deletion allele. (3) One of the supposedly supporting reads in the mate-pair library has the read-pairs overlapping each other (indicated by a green highlight in the middle of the read), despite an insert size of 6.3±1.2kb, which is highly unlikely.

